# Spleen Transcriptome Analysis of C57BL/6 Mouse in Response to Hot Water Extract from Spent Mushroom Substrate of *Ganoderma lucidum* and Cyclophosphamide

**DOI:** 10.1101/784108

**Authors:** Zehui Wang, Annan Wang, Jing Li, Zhen Liao, Lianyue Sun, Zhanxi Lin, Yanling Liu

**Author notes:** These authors contributed equally to this work (Zehui Wang, Annan Wang). These authors also contributed equally to this work (Zhen Liao, Lianyue Sun).

## Abstract

Previously, we have indicated that Hot Water Extract (HWE) from Spent Mushroom Substrate (SMS) of *Ganoderma lucidum* enhanced immune function of normal mice, and improved antioxidant activity and enhanced immune function of immunosuppression mice induced by cyclophosphamide (Cy). Here we performed the high throughput RNA sequencing strategy using Illumina HiSeq™ 2000 to characterize the spleen transcriptome from normal (CK1), HWE-treated (CK2), Cy-treated (CY) and both high dose HWE and Cy-treated mice (CH). From the RNA Sequencing, total mapped reads of map to Gene in CK1, CK2, CY and CH was 54 759 942, 54 678 926, 44 728 132 and 54 006 596, respectively. And gene expression was significantly different among CK1 and CK2, CY and CH. Compared with CK1, the gene expression of *Ugt1a6b* was down-regulated in CK2 after HWE treated. In addition, compared with CY, multiple tumor suppressor or tumorigenesis genes were down-regulated, such as *Cdkn1a*, *Cdkn1b*, *Mapk*10, *Vash1*, and *Tnc* and other genes in CK2 and CH. Taken together, our study highlighted the spleen transcriptome profiles of C57BL/6 mouse in response to HWE from SMS of *G. lucidum* and Cy, and indicated that HWE can improve the immune function of the mouse and accelerated the recovery of immunosuppression in Cy-treated mice.

## Introduction

*Ganoderma lucidum* named Lingzhi in Chinese, a precious traditional medicinal fungus in Asia, has been widely known for centuries as an immunomodulating agent, regulating many diseases, including gastric cancer [1], hypertension [2], arthritis [3], chronic hepatitis [4], diabetes [5], asthma [6], nephritis [7], arteriosclerosis [8], and immunological disorders [9]. The effectiveness of *G. lucidum* has been attributed to the polysaccharides fraction, which is responsible for the stimulation of immune system [10, 11]. The *G. lucidum* polysaccharides (GLP) are believed to trigger an indirect antitumor mechanism in which the host immune system is altered to target the tumor cells, meanwhile, it has been shown that GLP have the ability to induce both innate and adaptive immune response [12–16]. The polysaccharides from Chinese herbal medicine are regarded to involve in the activity regulation of immune cells and immune-related cells including T lymphocytes [17], B lymphocytes [18], macrophages [19], dendritic cells [20], and natural killer (NK) cells [21].

The main bioactive component of hot water extract (HWE) from Spent Mushroom Substrate (SMS) of *G. lucidum* is polysaccharides (15.79%). Interestingly, our previous studies have suggested that HWE can enhance the immune function in normal mice, shown that HWE can enhance markedly the spleen lymphocytes proliferation caused by ConA, delayed typed hypersensitivity induced by DNFB, on chicken RBC phagocytizing ability of peritoneal macrophages, the ability of charcoal particles clearance, improve the formation of antibody - producing cells and natural killer cell activity in mice [22], in accordance with the reported observation that GLP enhanced the function of immunological effector cells in immunosuppressed mice [23, 24].

It is well known that cyclophosphamide (Cy) is a crucial chemotherapeutic drug in tumor treatment, also induces numerous adverse effects, especially myelosuppression [25] and immunosuppression [26, 27]. Mice treated with Cy via intraperitoneal injection are usually used as an immunosuppression model [28]. In addition, HWE improves the recovery of suppressed immune function in mice induced by Cy shown by increasing the mouse spleen and thymus indices, and total anti-oxidative activity [29], and inhibiting the concentration of interleukin- 1β, interferon-γ, and tumor necrosis factor-α in the serum of immune-deficient mice [30]. However, the mechanisms underlying the immunomodulating effect of HWE remain uncertain.

Crucial immune organs, such as spleen and thymus as well as immune cells, including lymphocytes, macrophages, natural killer (NK) cells, play significant roles in anti-cancer effect [31, 32]. The spleen combines the innate and adaptive immune system in a uniquely organized way. The spleen is the largest filter of the blood in the body and unique among the lymphoid organs. Within the spleen, erythrocytes and iron are recycled, blood-borne pathogens are captured and destroyed, and both innate and adaptive immune responses can be mounted in response to circulating antigens [33].

The aim was to explore the spleen transcriptome with the effects of HWE in mice. Firstly, we established the immune-deficient mice model in C57 BL/6 mice induced by Cy, gene expression of the mice spleen were detected among the control group (CK1), the normal control group (CK2) treated with HWE, the Cy model group (CY) treated with Cy and the group (CH) treated with high dose HWE and Cy, and many differentially expressed genes (DEGs) were identified. Secondly, we focused the analysis of the spleen gene expression between CK1 vs CK2, CY vs CH, CY vs CK2, to emphasize and explain the effect of HWE in normal mice and in immune-deficient model mice. Among these DEGs, there were many genes related to immune system and cancer. In this study, the results may help to better understand that some genes and pathways are involved in regulating the immune response to HWE and Cy.

## Materials and Methods

### Hot Water Extract from Spent Mushroom Substrate of *G. lucidum*

*G. lucidum* was provided by the National Engineering Research Center of JUNCAO Technology (Fuzhou, China) and cultivated using JUNCAO technology, which was performed in 1986. Lin Zhanxi and his team discovered that JUNCAO species, such as *N. reynaudiana*, *S. anglica*, *M. floridulus* and *P. purpureum* are high-quality culture materials for *G. lucidum* cultivation. The most commonly JUNCAO used as culture media are: *Dicranopteris dichotoma* (Thunb.) Berhn 38%, *Miscanthus floridulus* 40%, wheat bran 20%, gypsum 2%. The moisture content of the medium is about 60%. Under certain suitable conditions, the cultivation of *G. lucidum* using JUNCAO technology can be performed in two months from germination to harvest.

After the harvest, Spent Mushroom Substrate (SMS) of *G. lucidum* were collected. HWE was obtained by boiling the SMS of *G. lucidum* in water for 4 h at the ratio 10:1 water to raw materials and then filtering the mixture. The residue was collected and extracted twice more with boiling water for 3 h at the ratio 8:1 water to raw materials. Then, the resulting supernatant was concentrated under vacuum conditions with the degree of vacuum at 0.09 MPa and the temperature at 55°C-65°C. HWE powders were produced by a spray-drying process. Ten kilograms of HWE were obtained from 100 kg of fresh SMSG. The HWE included 23.58% crude protein, 17.60% ash, 4.95% amino acid, and 2.0% crude fat. The content of polysaccharide in HWE was 15.79% as determined by high-performance liquid chromatography using the standard provided by National Engineering Research Center of JUNCAO Technology.

### Animals and experimental design

#### Ethics Statement

All the animal care and protocols were approved by the Animal Care Committee of Peking University Health Center (approval no. LA2015055). All experiments were conducted in accordance with the guidelines for the Care and Use of Laboratory Animals [34]. All procedures described below were conducted in accordance with the Inspection and Evaluation of Health Food [35]. Adequate measure were taken to minimize pain of experiment animals, and animal welfare was monitored twice daily by assessment of clinical conditions and weight change of mice.

#### Experimental treatments and sample collection

Sixty, pathogen-free, male C57BL/6 mice (7 weeks old, weighing 18-20 g) were purchased from Charles River Laboratories (Beijing, China), the animal license number for the mice is SCXK (Beijing) 2012-0001., China. The mice underwent one week of acclimation to the laboratory conditions prior to treatment. The mice were provided with standard chow and water *ad libitum* under the conditions at 24±1°C with a 12-h photoperiod.

Mice were randomly allocated into six groups (n = 10 each, Table 1), were intraperitoneally injected once a week (day1, day8, day 15, day 22) for 28 days of (1) control group (CK1), normal group (CK2) (100mg/kg normal saline), (2) Cy treatment groups (100 mg/kg Cy, which was purchase from Shanxi Pude Pharmaceutical CO., Ltd. (Batch No. 04120101, Shanxi, China)). Mice were weighted every day, and HWE was given to the mice of CK2, CL, CM and CH by oral administration every day, and the dose is 1g/kg BW, 0.5g/kg BW, 1g/kg BW and 2g/kg BW, respectively, while the mice of CK1 and CY were administrated with water by oral every day.

**Table 1.**
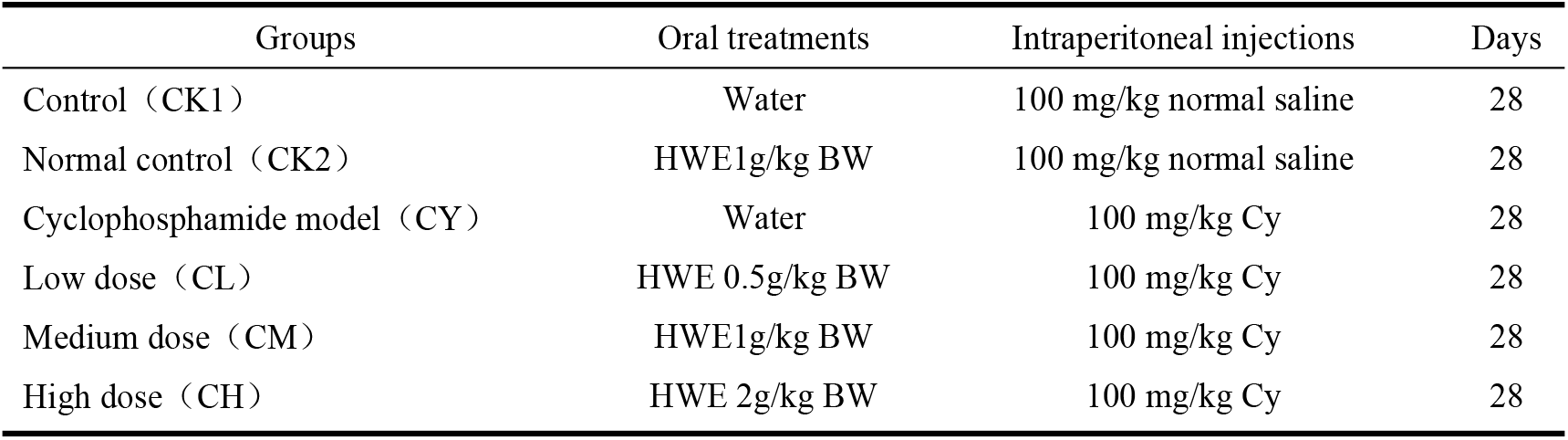
Establishment and the program of immunosuppression mice model.

| Groups | Oral treatments | Intraperitoneal injections | Days |
| --- | --- | --- | --- |
| Control (CK1) | Water | 100 mg/kg normal saline | 28 |
| Normal control (CK2) | HWE1g/kg BW | 100 mg/kg normal saline | 28 |
| Cyclophosphamide model (CY) | Water | 100 mg/kg Cy | 28 |
| Low dose (CL) | HWE 0.5g/kg BW | 100 mg/kg Cy | 28 |
| Medium dose (CM) | HWE1g/kg BW | 100 mg/kg Cy | 28 |
| High dose (CH) | HWE 2g/kg BW | 100 mg/kg Cy | 28 |

On the twenty-ninth day, mice were sacrificed by cervical dislocation. Spleen were immediately removed under sterile environment and stored in −80℃ for the RNA extraction.

### Gene expression analysis

#### RNA Extraction, cDNA library construction and Sequencing

The total RNA was isolated using the Trizol Kit (Thermo Fisher Scientific, USA) using standard methods and homogenized using a Tissue Lyser II (QIAGEN, Germany). Tissue with Trizol was extracted according to the manufacture instructions. Then the total RNA was treated with RNase-free DNase I (Takara Bio, Japan) for 30 min at 37°C to remove residual DNA. RNA quality was verified using a 2100 Bio analyzer (Agilent Technologies, Santa Clara, CA) and were also checked by RNase free agarose gel electrophoresis.

Next, Poly (A) mRNA was isolated using oligo-dT beads (Qiagen). All mRNA was broken into short fragments by adding fragmentation buffer. First-strand cDNA was generated using random hexamer-primed reverse transcription, followed by the synthesis of the second-strand cDNA using RNase H and DNA polymerase I. The cDNA fragments were purified using a QIA quick PCR extraction kit. These purified fragments were then washed with EB buffer for end reparation poly (A) addition and ligated to sequencing adapters. Following agarose gel electrophoresis and extraction of cDNA from gels, the cDNA fragments were purified and enriched by PCR to construct the final cDNA library. The cDNA library was sequenced on the Illumina sequencing platform (Illumina HiSeq™ 2000) using the paired-end technology by Gene De novo Co. (Guangzhou, China).A Perl program was written to select clean reads by removing low quality sequences (there were more than 50% bases with quality lower than 20 in one sequence), reads with more than 5% N bases (bases unknown) and reads containing adaptor sequences.

#### Reads alignment and Normalization of gene expression levels

Sequencing reads were mapped to reference sequence by the SOAP aligner/soap2 [36], a tool designed for short sequences alignment. Coverage of reads in one gene was used to calculate expression level of this gene. Using this method we obtained the expression levels of all genes detected.

Reads that could be uniquely mapped to a gene were used to calculate the expression level. The gene expression level was measured by the number of uniquely mapped reads per kilobase of exon region per million mappable reads (RPKM). The formula was defined as below:

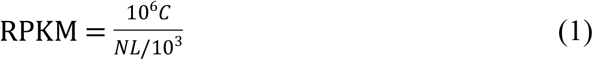

In which C was the number of reads uniquely mapped to the given gene; N was the number of reads uniquely mapped to all genes; L was the total length of exons from the given gene. For genes with more than one alternative transcript, the longest transcript was selected to calculate the RPKM. The RPKM method eliminates the influence of different gene lengths and sequencing discrepancies on the gene expression calculation. Therefore, the RPKM value can be directly used for comparing the differences in gene expression among samples. All expression data statistic and visualization was conduction with R package (http://www.r-project.org/).

#### Gene ontology analysis of differentially expressed genes (DEGs)

Gene Ontology (GO) is an international standardized gene functional classification system which offers a dynamic-updated controlled vocabulary and a strictly defined concept to comprehensively describe properties of genes and their products in any organism. GO has three ontologies: molecular function, cellular component and biological process. The basic unit of GO is GO-term. Every GO-term belongs to a type of ontology. All DEGs were blasted to GO database. We statistic it and show it in the result.

#### Differentially expressed genes (DEGs) and function enrichment analyses

After the expression level of each gene was calculated, differential expression analysis was conducted using edgeR [37]. The false discovery rate (FDR) was used to determine the threshold of the p value in multiple tests, and for the analysis, a threshold of the FDR≤0.01 and an absolute value of log2Ratio≥ 1 were used to judge the significance of the gene expression differences.

DEGs were used for GO and KEGG enrichment analyses according to a method similar to that described by Zhang [38]. Both GO terms and KEGG pathways with a Q-value ≤0.05 are significantly enriched in DEGs.

Gene function was annotated based on the following databases, Kyoto Encyclopedia of Genes and Genomes (KEGG), and Gene Ontology (GO). GO enrichment analysis of DEGs was implemented using the top GOR packages. KOBAS software was used to test the statistical enrichment of DEGs in KEGG pathways.

## Results

### RNA assessment and basic statistics of samples

The RNA quality and concentration of the spleen samples were shown in the Table 2. The total amount RNA ranged from 33μg to46 μg, and the OD 260/280 from 1.90 to 2.0. The results indicated that the RNA of all samples was integrated and stable for the RNA sequencing.

**Table 2.** Concentration and quality of the RNA samples.

| **Sample** | **Concentration(ng/mL)** | **A260/280** | **Total RNA (μg)** |
| --- | --- | --- | --- |
| **CK1-7** | 342 | 1.91 | 34.2 |
| **CK1-9** | 413 | 1.97 | 41.3 |
| **CK1-10** | 338 | 1.96 | 33.8 |
| **CK2-2** | 346 | 1.92 | 34.6 |
| **CK2-5** | 386 | 1.98 | 38.6 |
| **CK2-7** | 393 | 1.99 | 39.3 |
| **CY-4** | 456 | 1.92 | 45.6 |
| **CY-5** | 421 | 1.96 | 42.1 |
| **CY-7** | 439 | 1.97 | 43.9 |
| **CH3** | 402 | 1.95 | 40.2 |
| **CH6** | 442 | 1.97 | 44.2 |
| **CH8** | 458 | 1.96 | 45.8 |

From the RNA sequencing, total mapped reads of map to Gene in CK1, CK2, CY and CH was 54 759 942, 54 678 926, 44 728 132 and 54 006 596, respectively (Table 3). And the base composition of clean reads in CK1, CK2, CY and CH was shown in S1 Fig, while S2 Fig showed the random distribution of sequencing reads in assembled unigenes.

**Table 3.** Statistics of each sample CK1, CK2, CY and CH.

|  | **CK1** | **CK2** | **CY** | **CH** |
| --- | --- | --- | --- | --- |
| **Clean reads** | 54759942 | 54678926 | 54006596 | 54006596 |
| **Genome map Rate** | 85.61% | 82.17% | 82.84% | 80.37% |
| **Gene map Rate** | 73.93% | 78.20% | 74.12% | 80.81% |
| **Expressed Gene** | 16429 | 16325 | 16730 | 16240 |
| **Expressed Transcripts** | 21260 | 21176 | 21710 | 21029 |
| **Expressed Exon** | 187171 | 189588 | 195325 | 188638 |
| **Novel Transcripts** | 809 | 711 | 905 | 626 |
| **Extend Gene** | 2433 | 2170 | 2682 | 2069 |
| **Alternative Splicing** | 47047 | 48519 | 45940 | 47022 |
| **SNP** | 24410 | 24756 | 24341 | 24017 |

### Significantly differentially expressed genes

The results of differentially expressed genes indicated that (Fig 1), compared with CK1, there were 2,155 genes significantly down-regulated and 1,318 genes significantly up-regulated in CH. 216 genes were down-regulated and 500 genes were significantly up-regulated in CK2 compared with CK1. Compared with CK2, there were 1035 genes significantly down-regulated and 440 genes significantly up-regulated in CH.

**Fig 1.**
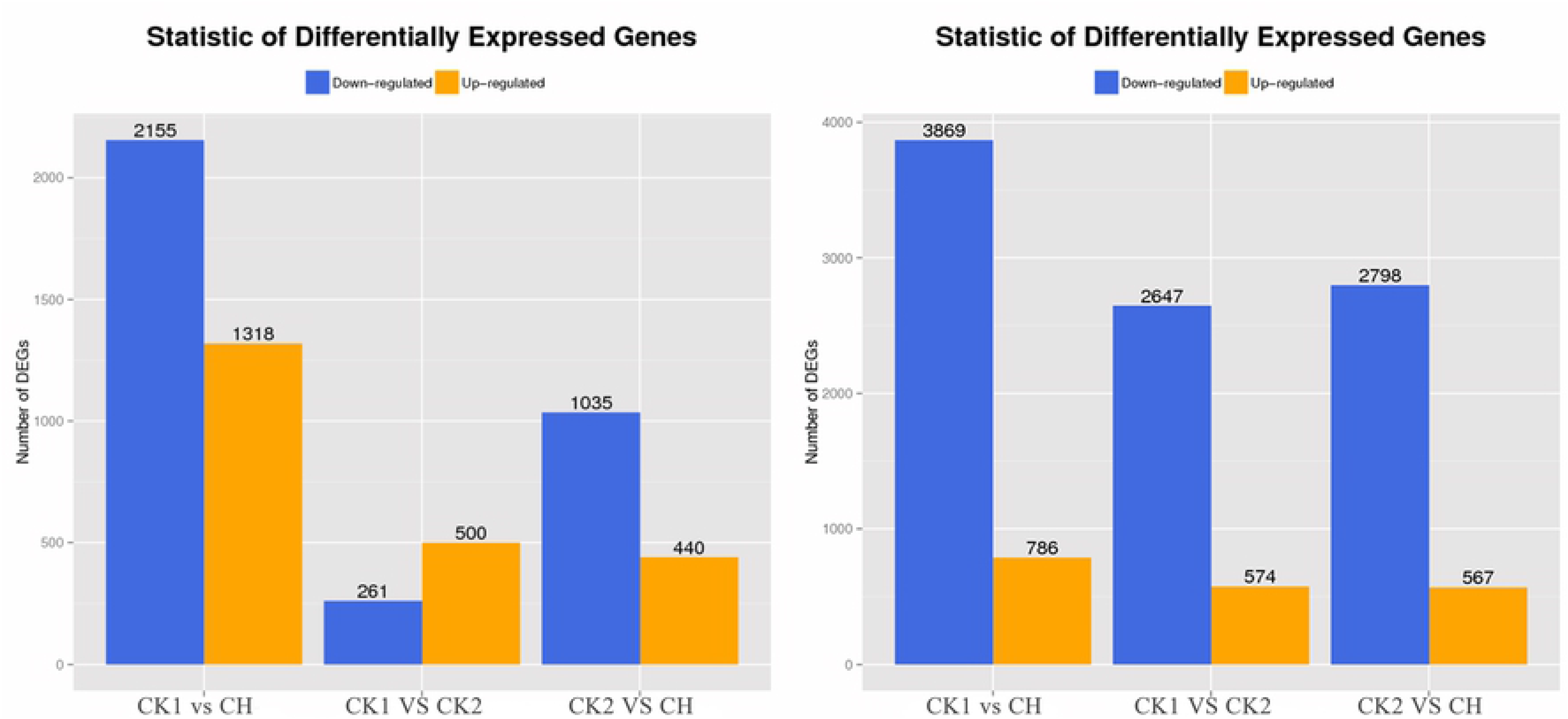
Statistical chart of gene expression with significant differences.

Compared with CY, there were 3869 genes significantly down-regulated while 786 genes were significantly up-regulated in CH, 2647 genes were significantly down-regulated and 574 genes were significantly up-regulated in CK1, there were 2798 genes significantly down-regulated and 567 genes significantly up-regulated in CK2.

The scatter and volcano diagram (S3 Fig) results showed significantly differentially expressed genes (DEGs) in the above comparisons.

### Gene ontology analysis of differentially expressed genes

To generate the functional distribution of the differentially expressed genes (DEGs), a GO analysis of these genes was performed. For the comparison of CK1 vs CK2 (Fig 2), CY vs CH (Fig 3), CY vs CK2 (Fig 4), the differentially expressed genes were classified into three major functional categories. Among them, the largest numbers of DEGs were clustered in cellular component-561genes for CK1 vs CK2, 3931 genes for CY vs CH, 2839 genes for CY vs CK2, followed by biological process-545 genes for CK1 vs CK2, 3751 genes for CY vs CH, 2699 genes for CY vs CK2, and molecular function-541 genes for CK1 vs CK2, 3688 genes for CY vs CH, 2645 genes for CY vs CK2.

**Fig 2.**
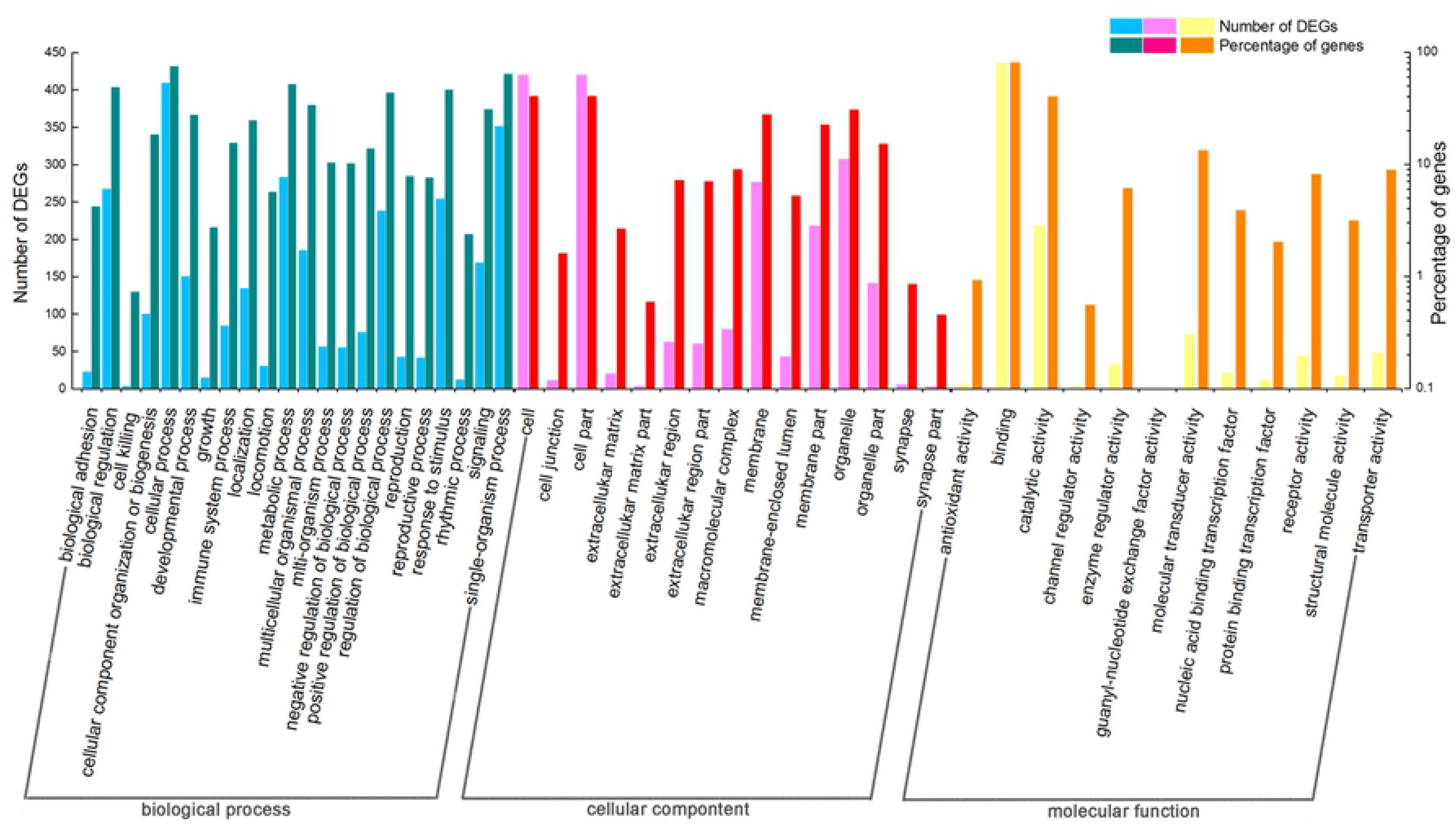
GO functional classification analysis in gene expression with significant differences among CK1 vs CK2.

**Fig 3.**
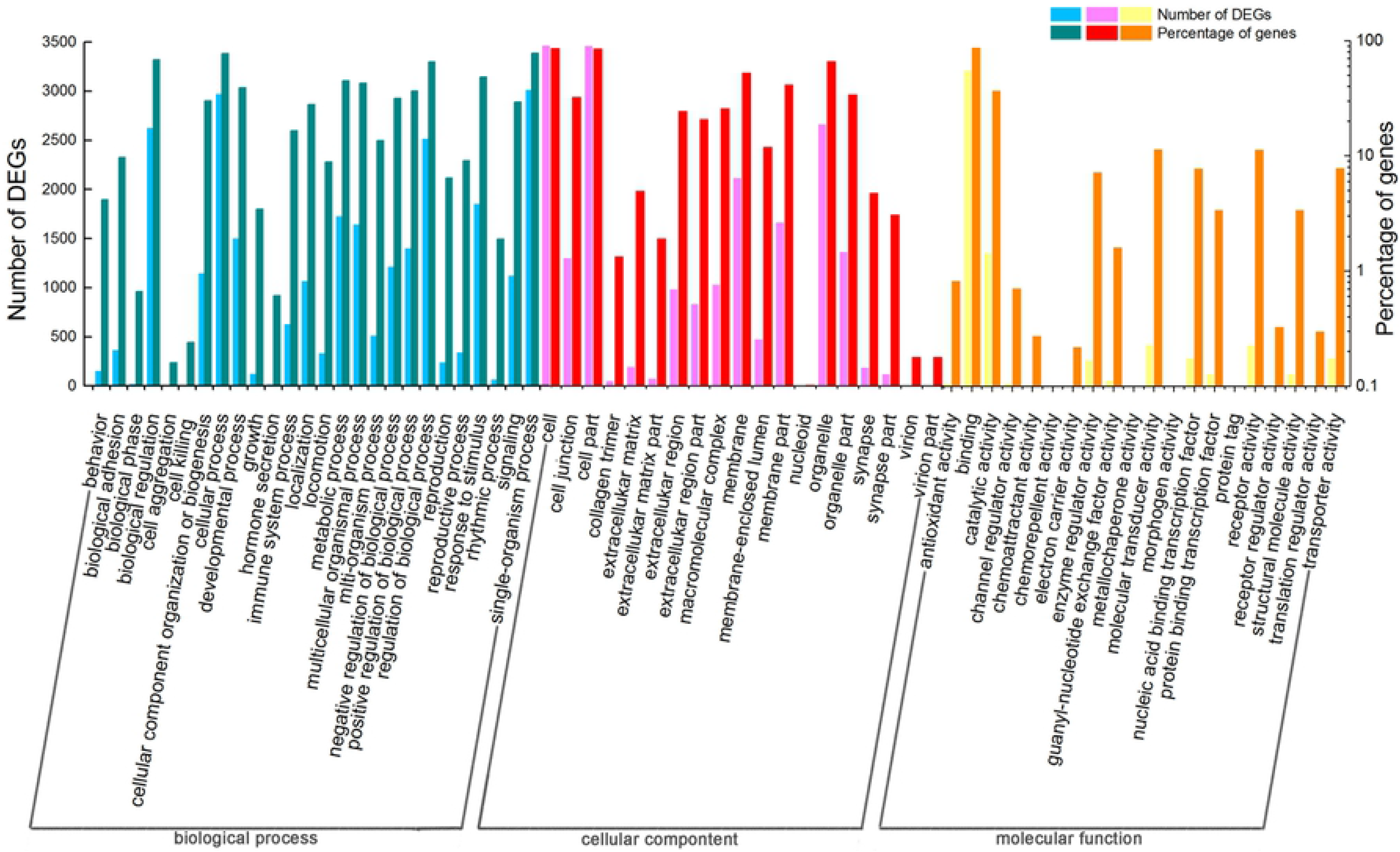
GO functional classification analysis in gene expression with significant differences among CY vs CH.

**Fig 4.**
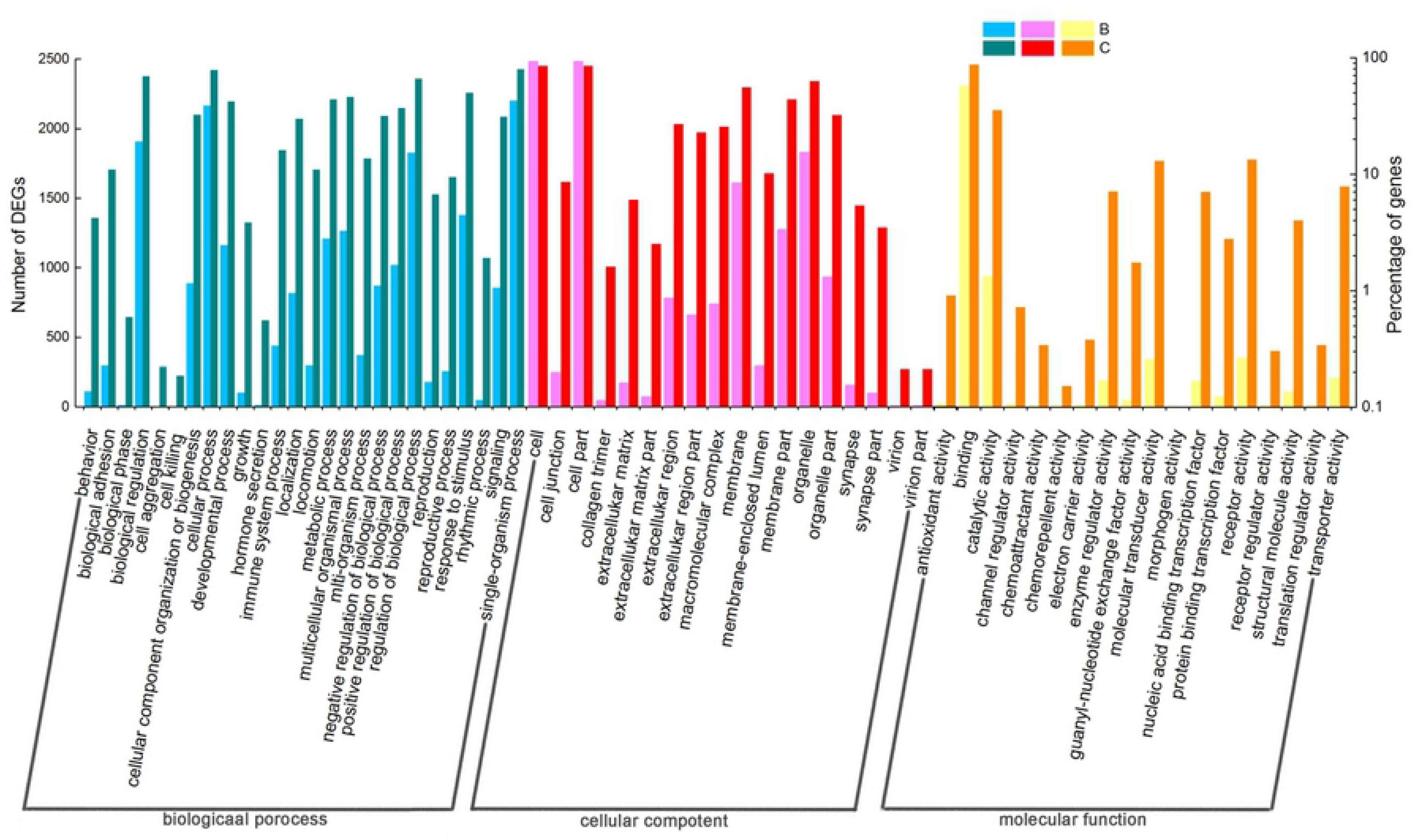
GO functional classification analysis in gene expression with significant differences among CY vs CK2.

### Pathway significant enrichment analysis of gene expression with significant differences

Additionally, we conducted a pathway significant enrichment analysis to generate overviews of the functions and interactions of these DEGs. To further understand the differentially expressed genes under the treatments of HWE and cyclophosphamide, the followings are the results of the comparison between CK1 (control) and CK2 (with HWE), CY (with cyclophosphamide) and CH (with HWE and cyclophosphamide), CY (with cyclophosphamide) and CK2 (with HWE).

#### CK1 vs CK2

The important pathways between CK1 and CK2, obtained from pathway enrichment for gene expression with significant differences, included Staphylococus aureus inferction, Prophyrim and chlorophyll metabolism, Phagosome, Malaria, Hypertrophic cardiomyopathy, Dilated cardiomyopathy, Glycerolipid metabolism and so on (Fig 5). Table 4 showed some differential expression of gene associated with immune according to the results of top 20 statistics of pathway enrichment for gene expression with significant differences between CK1 and CK2.

**Fig 5.**
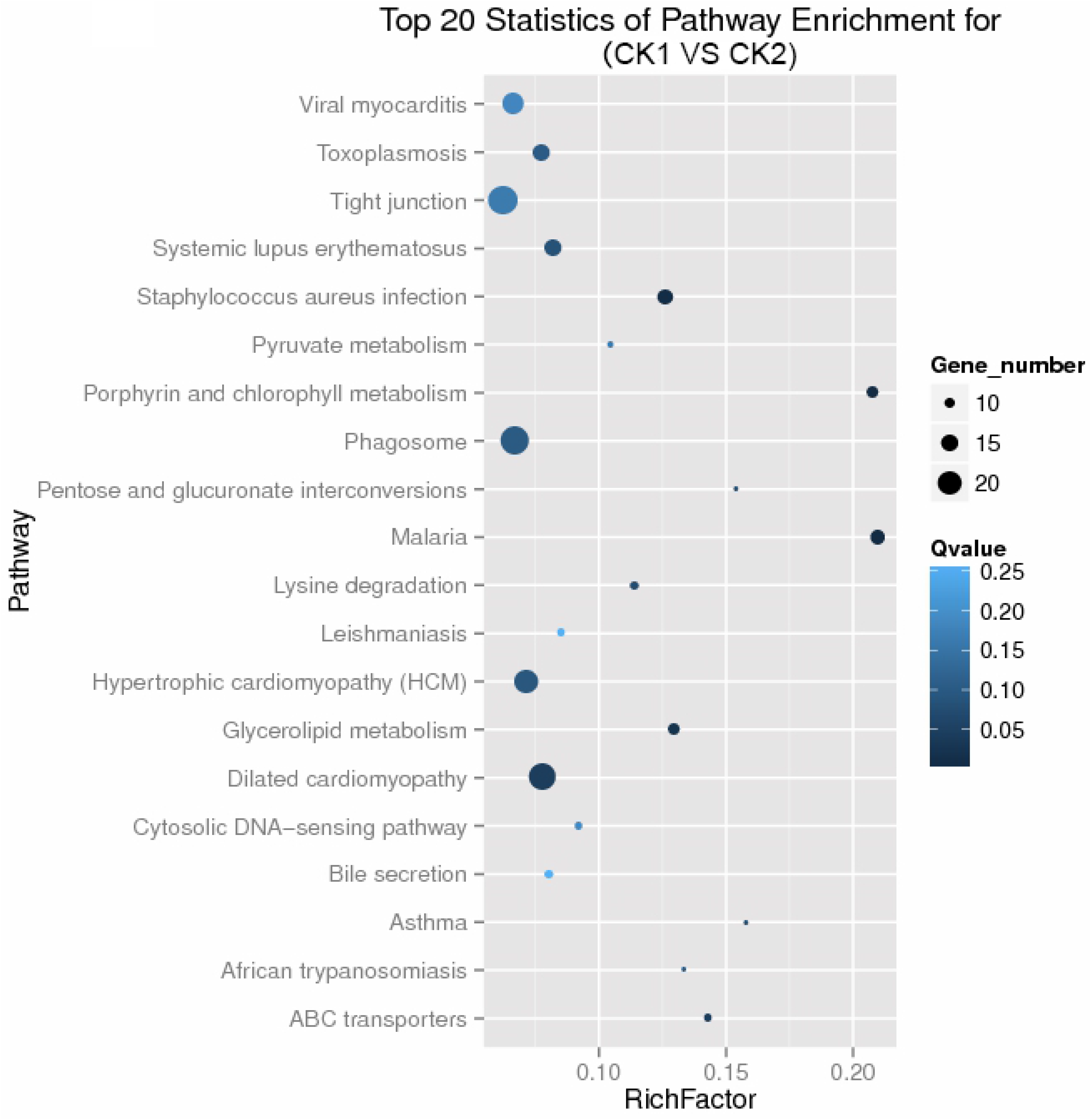
Top20 Statistics of Pathway Enrichment in differentially expressed genes for CK1 vs CK2.

**Table 4.**
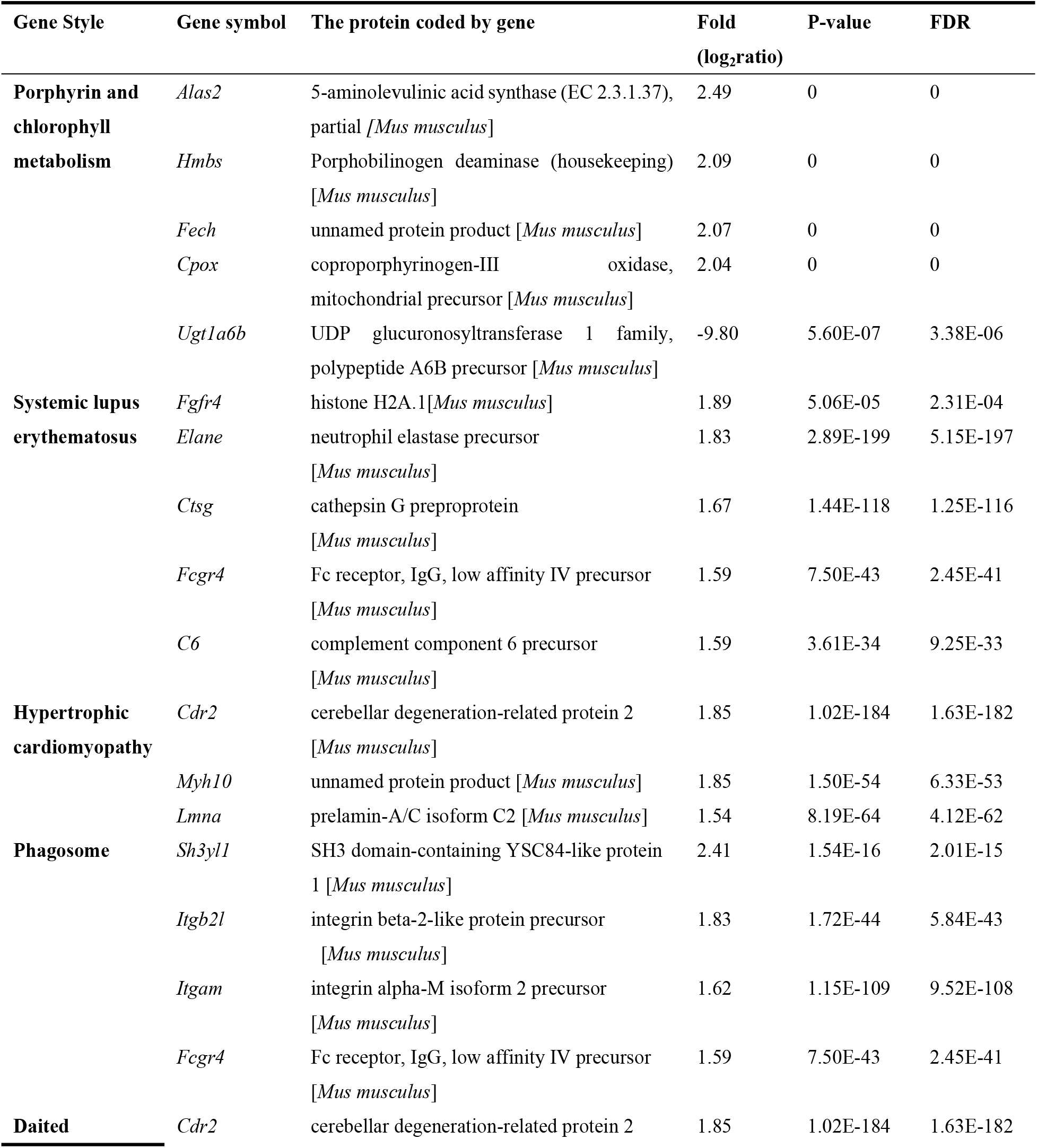
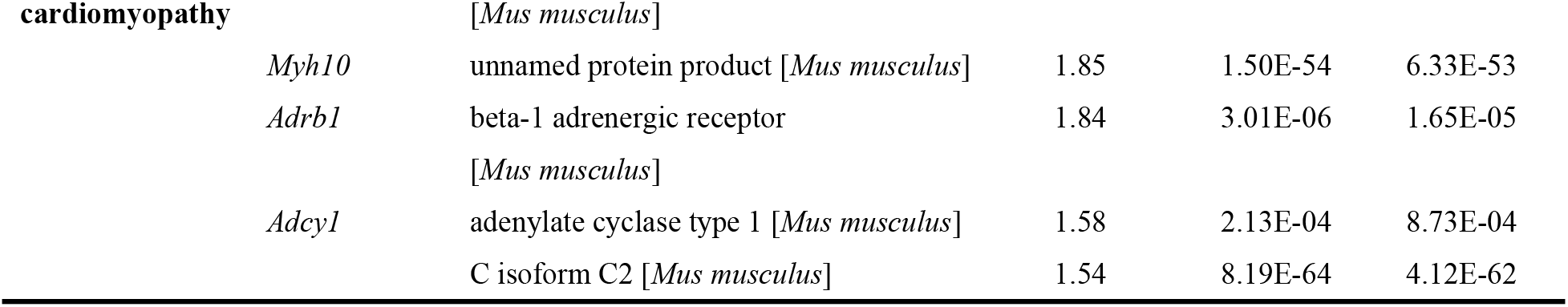
Differential expression of gene associated with immune between CK1 and CK2.

The results showed that, compared with CK1, the gene expression of *Alas2* in CK2, encoding 5-aminolevulinic acid synthase, an aminolevulinic acid synthase gene, was up-regulated, 2.49 times of that in CK1, *Ugt1a6b* (UDP glucuronosyltransferase 1 family, polypeptide A6B precursor) was down-regulated.

#### CY vs CH

The important pathways in Fig 6, which showed the results of top 20 statistics of pathway enrichment for gene expression with significant differences between CY and CH, referred to Pathway in cancer, T-cell receptor signaling pathway, MAPK signaling pathway, HTLV-1 interaction, Focal adhesion, Cells adhesion molecules, ECM-receptor interaction, Toxoplasmosis, Axon guidance and so on. Table 5 showed some differential expression of gene associated with immune according to the results of top 20 statistics of pathway enrichment for gene expression with significant differences between CY and CH.

**Fig 6.**
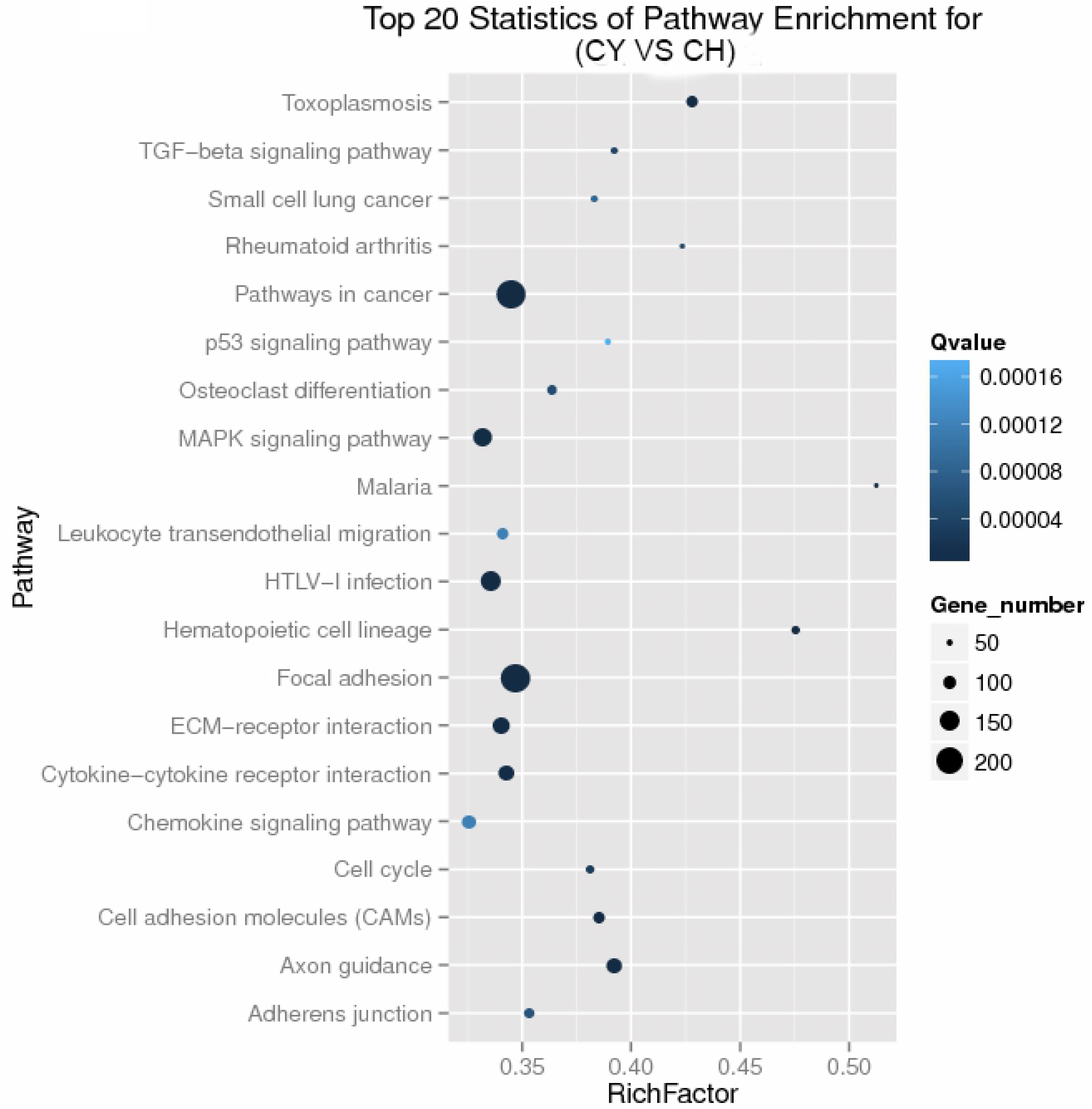
Top20 Statistics of Pathway Enrichment in differentially expressed genes for CY vs CH.

**Table 5.**
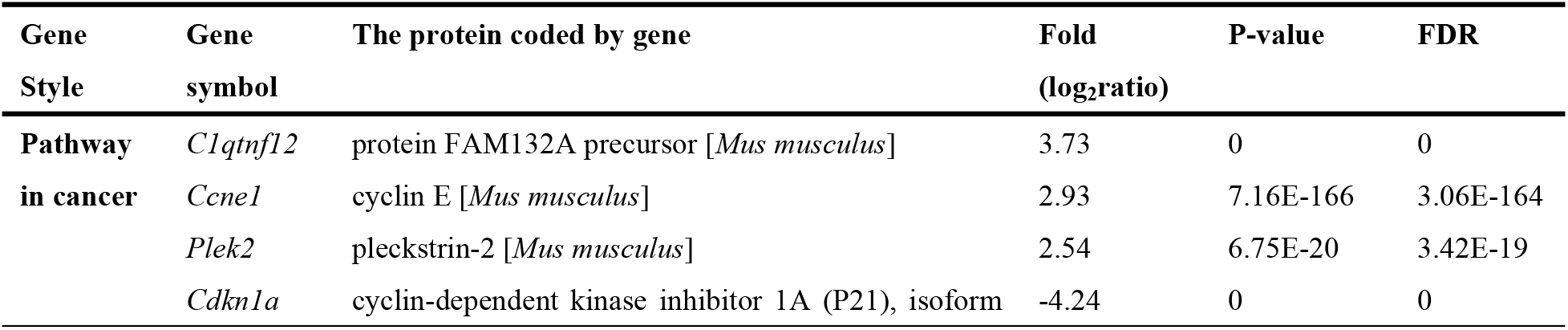
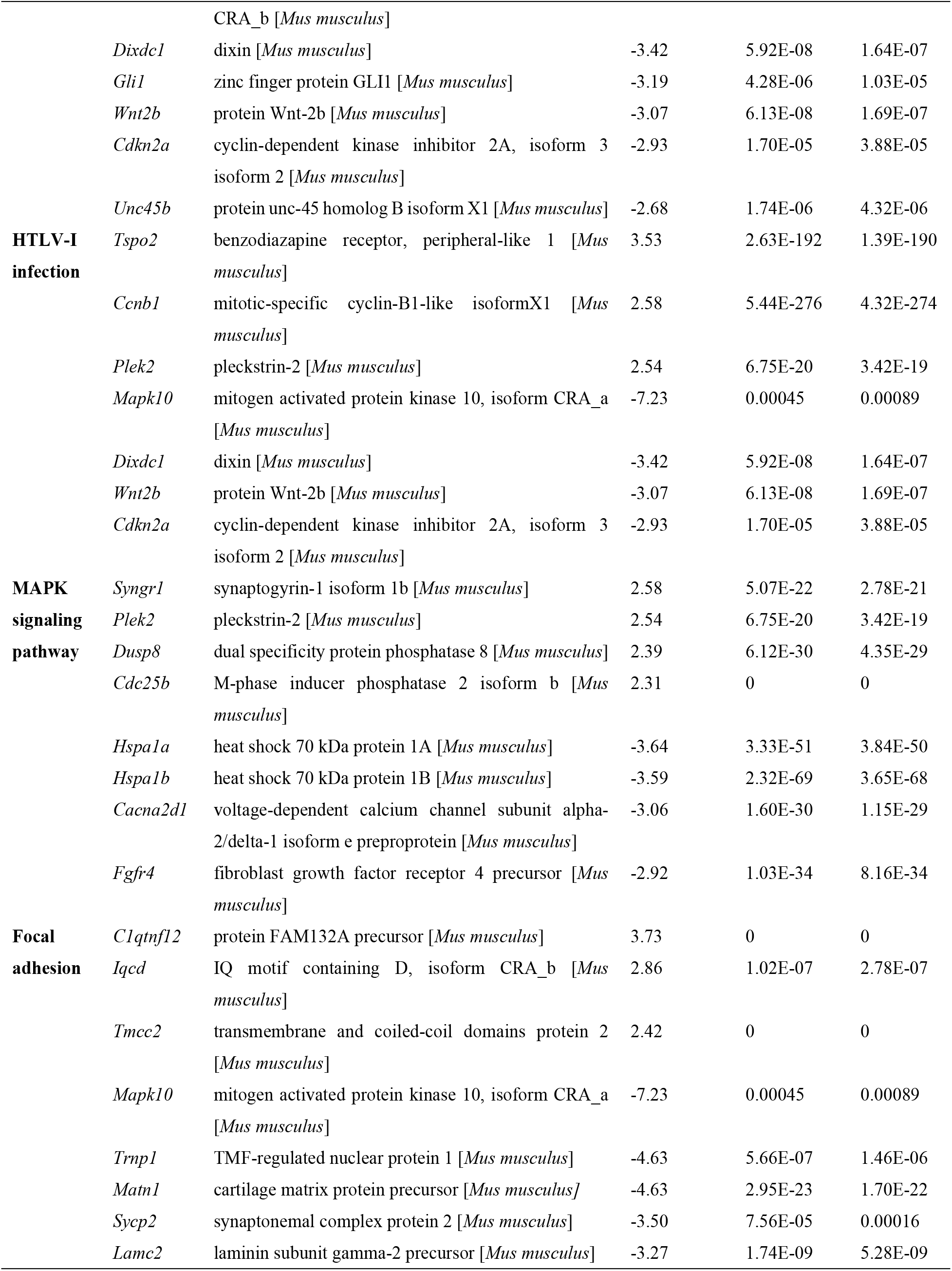
Differential expression of gene associated with immune between CY and CH.

The expression of gene *Ccne1* (cyclin E) in the high-dose group of mouse spleen were 2.93 times expression in the CY model group. Compared with CY, the gene *Cdkn1a* of the spleen was down-regulated in CH, and the expression of *Cdkn1a* in CY was 4.24 times higher than that in CH. The gene expression of *Mapk10* (mitogen activated protein kinase 10, isoform CRA), *Trnp1* (TMF-regulated nuclear protein 1) and *Matn1* (cartilage matrix protein precursor) in CH was 7.23 times, 4.63 times and 4.63 times higher than that in CY, respectively.

#### CY and CK2

The important pathways in Fig 7, which showed the results of top 20 statistics of pathway enrichment for gene expression with significant differences between CY and CK2, referred to Regulation of actin cytoskeleton, Pathway in cancer, Focal adhesion, Cytokine-Cytokine receptor interaction, Dilated cardiomyopathy, Cells adhesion molecules (CAMs), ECM-receptor interaction, Axon guidance, Amoebiasis and so on. Table 6 showed some differential expression of gene associated with immune according to the results of top 20 statistics of pathway enrichment for gene expression with significant differences between CY and CK2.

**Fig 7.**
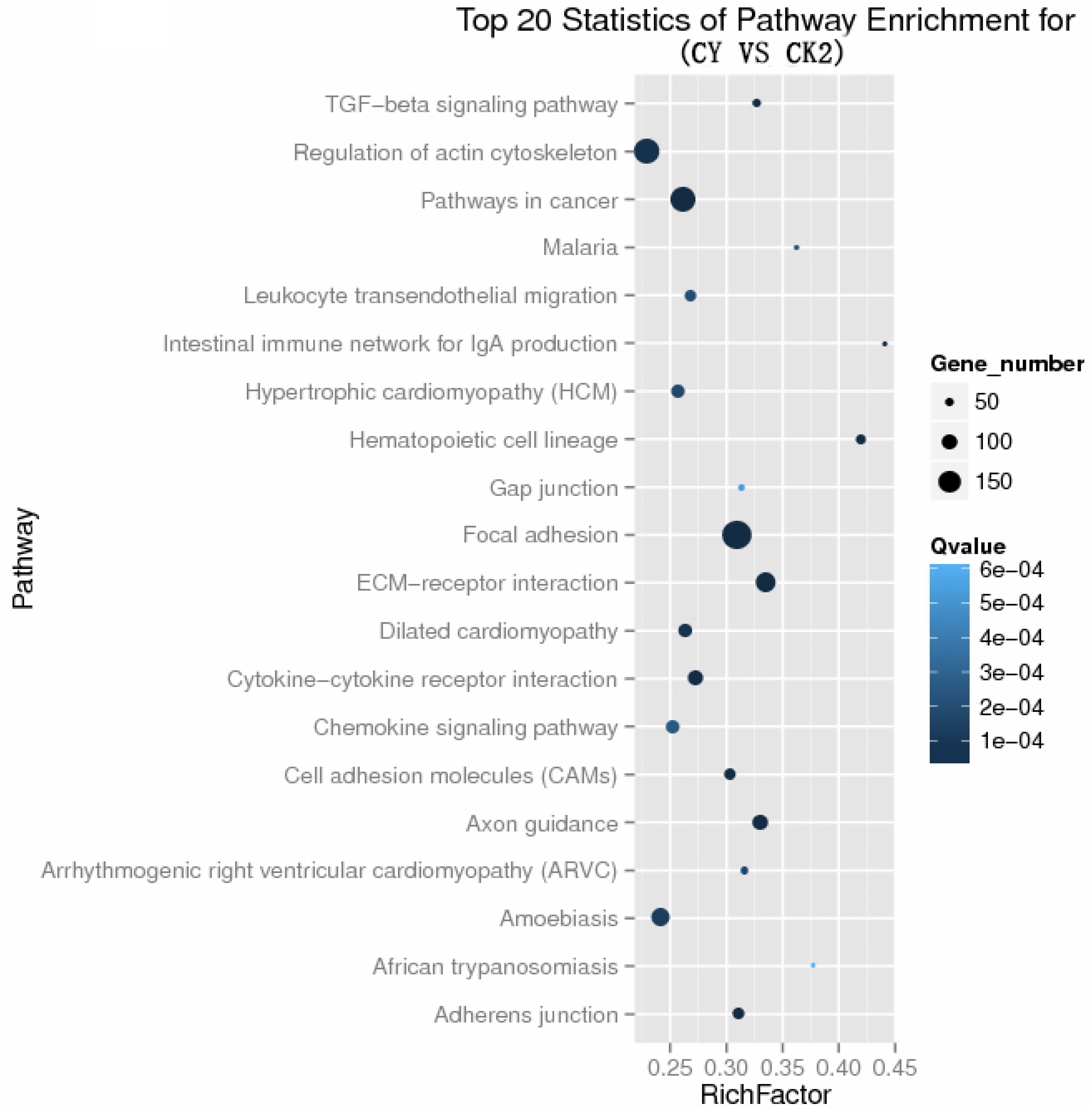
Top20 Statistics of Pathway Enrichment in differentially expressed genes for CY vs CK2.

**Table 6.**
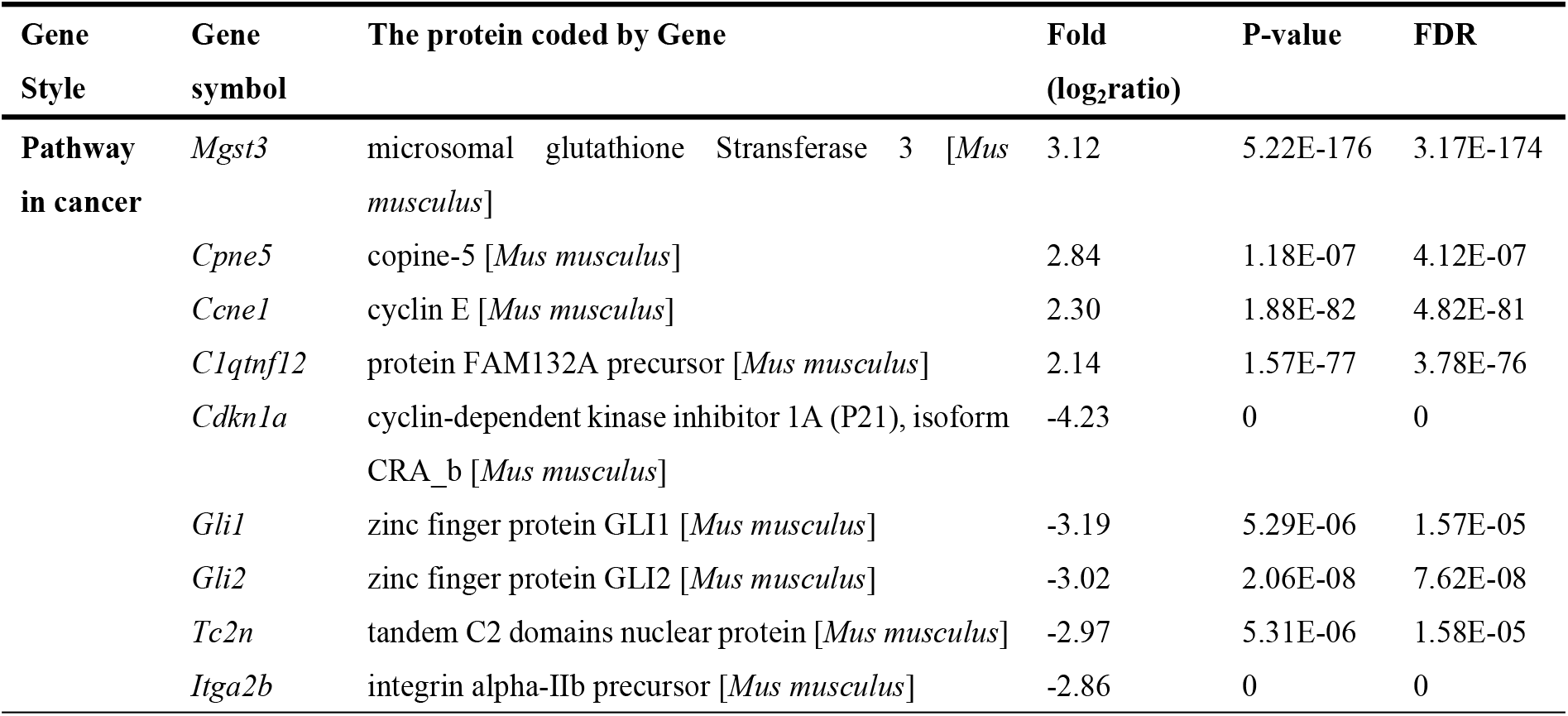
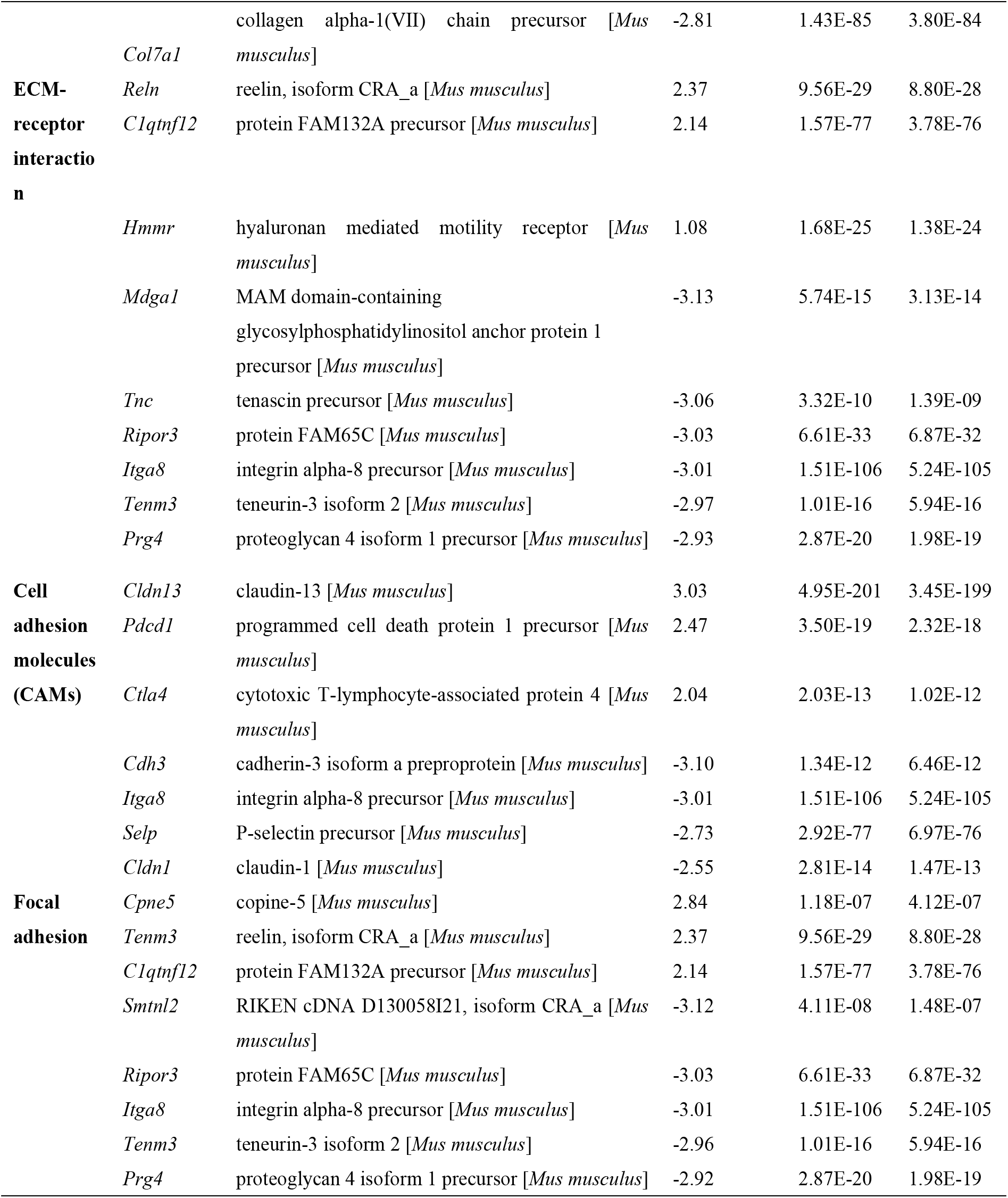
Differential expression of gene associated with immune between CY and CK2.

The expression level of gene *Tnc* in the CY model group was 3.06 times that of the normal group. After injection of cyclophosphamide, the mouse body was injured and the protein encoded by the *Tnc* gene was increased. The results showed that the expression level of gene *Gli2* in the CY model group was 3.02 times that of the normal control group, the expression of *Ctla4* in the normal control group was 2.04 times that of the model group. Genes such as *Mgst3*, *Cpne5*, *Reln*, *C1qtnf12*, *Hmmr*, *Pdcd1* were up-regulated, and most genes, such as *Gli1*, *GLli2*, *Tnc*, *Itga8*, *Mdga1*, *Tenm3*, *Prg4*, *Smtnl2*, *Ripor3* were down-regulated.

## Discussion

Notably, *G. lucidum* polysaccharide (GLP) is the major biologically active component in *G. lucidum* [39], which has anti-inflammatory and immunomodulatory effects. HWE, including 15.97% polysaccharide, is regarded as a functional feed additive, shown by improving milk yield, milk quality, hematology parameters, and enhancing immunity and antioxidant capacity in dairy cows [40, 41]. Our primary studies revealed that HWE enhances murine immune function [22], also improves the recovery of suppressed immune function in mice induced by Cy [29, 30]. However, it has yet to be determined whether HWE will affect the expression of genes related to immune-related pathways. In this study, using the immunosuppression model induced by Cy, differentially expressed genes among the treatments were identified using the high throughput sequencing. It was found that different pathways were enriched in different comparison of CK1 vs CK2, CY vs CH, CY vs CK2. The significantly enriched KEGG pathway not only included those related to immune system (human disease), but also those involved in cellular processes, organismal systems, as well as those that related to environmental information processing. These data indicated that when treated with HWE, amount of gene expression with or without cyclophosphamide in mice were affected.

### CK1 vs CK2

The results showed that, compared with Control group (CK1), 261 genes significantly were down-regulated meanwhile 500 genes up-regulated in Normal group (CK2).The significantly enriched KEGG pathways in the comparison of CK1 vs CK2 mainly belong to human disease, such as systemic lupus erythematosus, hypertrophic cardiomyopathy, dilated cardiomyopathy, viral myocarditis, toxoplasmosis, staphylococcus aureus infection, asthma, African trypanosomiasis. In addition, the main pathways of differential gene expression are also concentrated in pyruvate metabolism, pentose and glucose conversion, lysine degradation, glyceride metabolism and other pathways related to the body’s own metabolism.

The gene expression of *Ugt1a6b* was down-regulated in CK2 compared with CK1, and elevated by greater than 9.8 fold in the spleen of CK1, in accordance with that *Ugt1a6b* transcripts were elevated by greater than 3.0 fold in diabetic animals [42]. Normally, the immune function of the diabetic animals are lower than the normal animals. *Ugt1a6* catalyzes simple phenolic glucuronidation, and the neurotransmitter serotonin (5-hydroxytryptamine) is a typical endogenous substrate [43, 44]. There was few study which reported the Ugt1a6b gene expression in animals which are treated HWE or GLP. Intriguingly, there was no obvious immune-related pathways of top 20 enrichment analysis between CK1 and CH like T cell receptor signaling pathway, NK cytotoxicity and cytokine interaction. We speculated that HWE has a greater effect on the immune function of mice.

### CY vs CH

The differentially expressed genes results in the comparison of CY vs CH showed that, 3869 genes significantly were down-regulated, while 786 genes were up-regulated in high dose group (CH) compared with cyclophosphamide model group (CY). The expression of gene *Ccne1* in the high-dose group of mouse spleen were 2.93 times expression in the CY model group. Nakayama reported that *Ccne1* amplification had important therapeutic implications for patients with endometrial endometrioid carcinoma [45], and high *Ccne1* gene expression is a significant and independent predictor for prolonged overall survival in International Federation of Obstetrics and Gynecology (FIGO) II–IV epithelial ovarian cancer patients [46]. Furthermore, Cyclin E is associated with the development and progression of a variety of tumors, as Cyclin E is often deregulated in malignant cells, and this deregulation is often associated with tumor aggressiveness and poor prognosis [47]. In the present study, under the same injection times and the same dose of cyclophosphamide, the expression of Cyclin E and Cyclin A2 in the high-dose group was increased with the effect of HWE compared with CY model, we speculated that HWE did regulation and prognosis against the adverse reactions and damage caused by cyclophosphamide.

In pathway in cancer maps, p21 and p27 are proteins encoded by the genes *Cdkn1a* and *Cdkn1b*, respectively. The proteins p21Cip1 and p27Cip1 are both cip and kip families of cyclin-dependent kinase inhibitory proteins, which are both recognized as tumor suppressor. Deletion of p21Cip1 promotes tumorigenesis and tumor diversity in mice [48]. Watanabe et al found that *Cdkn1a* was specifically downregulated in adult T-cell leukemia/lymphoma cells compared with CD4^+^ T lymphocytes, while *Cdkn1a* was upregulated in the Human T-lymphotropic virus 1-infected cell lines[49]. Interestingly, our results found that the expression of *Cdkn1a* was upregulated in CY model group, 4.24 times higher than that in the high-dose group. Furthermore, Human T-lymphotropic virus 1-infection appeared in the significantly enriched pathway of top20, we speculated that it probably was related to Cy. Cy is a major constituent of cancer chemotherapy agent and widely used in the treatment of various types of cancer [50], meanwhile, immunosuppression induced by Cy increases incidence of secondary infections and mortality [51]. Hence, HWE may reduce the incidence of secondary infections and mortality caused by Cy and improve the immune function in Cy-treated mice.

### CY vs CK2

The gene expression with significant differences in the comparison of CY vs CK2 showed that 2798 genes were significantly down-regulated while 567 genes were up-regulated in CK2, compared with CY.

The protein encoded by gene *Tnc* is a glycoprotein, Tenascin C, which is an adhesion-regulating protein that can produce different functions in the same cell type and is dysregulated in pathological conditions such as inflammation, infection or tumor [52]. In this case, the expression level of gene *Tnc* in the CY model group was 3.06 times that of the normal group. Larger amounts are expressed or under pathological conditions such as inflammation, infection and tumorigenesis, during development, as well as in adults during tissue repair and remodeling, neovascularization, wound healing, and tumor genesis [53–58]. Since *Tnc* plays an important role in the degeneration of articular cartilage, and promotes cartilage repair, also strongly suggest that *Tnc* promotes chondrogenesis and cartilage repair in damaged and degenerated cartilage [59]. After injection of cyclophosphamide, we speculated that the mouse body was injured and the protein encoded by the *Tnc* gene was increased compared with the normal group treated with HWE.

The gene *Ctla4* encodes a cytotoxic T lymphocyte-associated protein 4, which acts as an immunoassay to down-regulate the immune system. *Ctla4* plays an important role in the regulation of T cells and their function, increasing expression of *Ctla4* by activating by T cell receptors and CD28, inhibiting transmitting signal to the T cells [60–63]. In this case, the expression of *Ctla4* in the normal control group was 2.04 times that of the model group. Conversely, the expression level of gene *Gli2* in the CY model group was 3.02 times that of the normal control group. Importantly, abnormal activation of the gene *Gli2* can lead to the occurrence of various malignant tumors such as pancreatic cancer, colon cancer, basal cell carcinoma, prostate cancer, neuroblastoma, and gastric cancer [64].

## Conclusion

In this study, using the cyclophosphamide-treated mice, and by the high throughput RNA sequencing verification, we have found that after HWE and cyclophosphamide treated, gene expression in mouse spleen, including many immune-related genes and many other genes. Taken together, the results indicated that the HWE can improve the immune function of the mouse and accelerated the recovery of immunosuppression in cyclophosphamide-treated mice.

## Acknowledgments

We thank Zhibin Lin, Weidong Li, Baoxue Yang (Peking University) for technical assistance and reagents.

## Supporting information

**S1 Fig. Base composition of clean reads in CK1, CK2, CY and CH** A: CK1; B: CK2; C: CY; D: CH. On the X axis of CK1, CK2 and CH, position 1-90bp represents read 1, and 90-180bp represents read 2. On the X axis of CY, position 1-100bp represents read 1, and 100-200 bp represents read 2. A curve nearly be overlapped with T curve while G curve overlapped with C curve. Red, green, blue, pink line each generation of each position A, T, C, G bases of proportion, Under the condition of base composition balance, A, T curve overlapping, G, C curve overlapping. If abnormal condition in the sequencing, base composition may not balance. The curve of the light blue represent each position are measured by the proportion of the base.

**S2 Fig. Random distribution of sequencing reads in assembled unigenes.** X-axis represents relative position of sequencing reads in the assembled sequences. Orientation of assembled unigenes is from the 5’ end to the 3’ end. Y-axis indicates number of reads.

**S3 Fig. Chart for differential expression:** X-axis and Y-axis presentation two samples log2value of expression, red(Up) and green(down) dot mean the gene has significant difference(FDR <=0.001,2 fold diffrence),and the black dot means no significant difference.

